# CaMKII induces an autophagy-dependent anabolic response in Articular Chondrocytes

**DOI:** 10.1101/2024.08.04.606243

**Authors:** Nicholas James Day, Angshumi Dutta, Cintia Scucuglia Heluany, Vipin Asopa, David Sochart, Barbara Fielding, Giovanna Nalesso

## Abstract

**Objective:** The objective of this study was to elucidate the role of Calcium calmodulin-dependent Kinase II (CaMKII) in articular chondrocytes and its involvement in osteoarthritis (OA) pathogenesis. By performing gain and loss of function experiments, the research aimed to determine how CaMKII modulates chondrocyte metabolism, anabolic and catabolic processes, hypertrophic differentiation, and autophagy within the articular cartilage.

**Design:** Articular cartilage was harvested from patients undergoing joint replacement surgery for OA, and adult human articular chondrocytes (AHACs) were isolated and cultured. Recombinant adenoviruses were used to overexpress a constitutively active form of CaMKIIγ (AdCaMKII) or inhibit CaMKII activity (AdAIP). Various assays, including RT-PCR analysis, alcian blue staining of Micromass cultures, immunofluorescence, and Western blotting, were performed to assess the effects of CaMKII modulation on chondrocyte function.

**Results:** Overexpression of activated CaMKIIγ promoted anabolism, evidenced by increased expression of SOX9, COL2A1, and ACAN, and decreased MMP-13 levels. It also enhanced proteoglycan content in AHAC micromass cultures. Furthermore, CaMKII counteracted the catabolic effects of IL-1β and preserved proteoglycan content. We also observed decreased chondrocyte proliferation and increased synthesis of hypertrophic marker Type X Collagen. CaMKII activation was found to induce autophagy, as indicated by increased phosphorylation of Beclin1 and decreased p62 expression. The anabolic effects of CaMKII were dependent on autophagy, as inhibition of autophagy with Bafilomycin prevented the CaMKII-induced increase in glycosaminoglycan content.

**Conclusions:** CaMKII plays a significant role in modulating chondrocyte metabolism and maintaining cartilage homeostasis. It promotes anabolic processes, counteracts catabolic stimuli, and induces autophagy in articular chondrocytes. However, it also promotes hypertrophic differentiation, highlighting the complexity of CaMKII-mediated signalling in cartilage. Understanding these pathways could lead to new therapeutic strategies that leverage CaMKII’s anabolic potential while mitigating its pro-degenerative effects.

## 1 Introduction

During osteoarthritis (OA), the balance between anabolism and catabolism within the articular cartilage (AC) is lost, leading to the breakdown of the Extracellular Matrix (ECM) and decline in the biomechanical properties of the tissue (1). OA pathogenesis is complex, and deregulation of several signalling cascades has been linked to the degeneration of the AC during disease progression.

We have previously shown that the activity of Calcium Calmodulin Dependant Kinase II (CaMKII) increases in the AC in OA (2), suggesting an important homeostatic function for this kinase within the tissue (2–5).

CaMKII is an intracellular serine-threonine kinase, which is expressed in mammals in four different isoforms: α, β, γ and δ (6). CaMKII has been extensively investigated in the brain for its involvement in regulating memory formation and synaptic plasticity (7). Within the AC, CaMKIIγ and δ are the two most abundant isoforms. While CaMKIIδ is expressed throughout the entire joint, CaMKIIγ is selectively expressed within the AC in mice (2). CaMKII activity has been associated with chondrocyte maturation and hypertrophy during bone development (4,8). Its role in the AC has only been partially characterised. Results so far suggest a pro-anabolic role for CaMKII as its pharmacological inhibition in a murine model of OA exacerbated cartilage damage and bone remodelling (2).

To increase our understanding of the role of CaMKII in articular chondrocytes and OA, here we performed gain and loss of function experiments aimed to better understand the function of this kinase in the articular chondrocytes. Our results point to CaMKII as an important modulator of chondrocyte metabolism at the intersection between multiple modulatory pathways within these cells.

## 2 Materials and Methods

### 2.1 Cartilage harvest and chondrocyte isolation

Adult human articular cartilage was obtained from patients who underwent joint replacement for OA of either the hip or knee after obtaining informed consent. Samples from 26 individual male and female donors were used during this study (1 patient< 50 year old, 2 patient > 80 year old, all the others in between 50 to 80 year old). The cartilage samples were provided by Prof D Sochart and Dr Vipin Asopa (SWLEOC, Epsom General Hospital, Surrey, UK). All procedures were approved by the HRA and Health and Care Research Wales (ethics approval REC N. 19/LO/0742). Full thickness explants were taken for histological scoring, and adult human articular chondrocytes (AHACs) were expanded and cultured as previously described (9). All experiments were carried out with either freshly isolated (FI) or passage 0 (P0) cells, from donors with a Mankin score of <4 (10).

Primary bovine chondrocytes were obtained from the metacarpophalangeal (MCP) and metatarsophalangeal (MTP) joints of domestic bovids purchased and collected from a local abattoir within 24h of slaughter. Since tissues and cells were collected from cows euthanized in a licensed abattoir and not specifically for this study, the Ethical Committee of the University of Surrey determined that no ethical approval was necessary. The cartilage was sliced and washed twice in DMEM/F12 + GlutaMAX (1:1) (1X) (Gibco) containing 10% FBS (Gibco Invitrogen, Carlsbad, CA, USA), 1mM sodium pyruvate and 2% antibiotic/antimycotic solution (Thermo Fisher, Waltham, MA, USA). Cartilage was then digested for 30 minutes at 37°C in 1 mg/ml pronase (Roche, Basel, Switzerland) and then overnight at 37°C in 1 mg/ml collagenase P (Roche, Basel, Switzerland) prepared in 10%CM (same composition as above, with 1% antibiotic/antimycotic solution) under agitation. Isolated chondrocytes were recovered were then seeded in complete media at 2 × 104/cm2 and cultured at 37°C containing 5% CO2. All experiments were conducted using freshly isolated, cells from the first passage (P0) or cells from the second passage (P1).

### 2.2 Adenoviral Transduction

To modulate CaMKII activity, we utilized recombinant adenoviruses. One adenovirus (AdCaMKII) overexpressed a constitutively active form of CaMKIIγ with a T287D mutation (11). Another adenovirus (AdAIP) expressed the Autocamtide-2 inhibitory peptide, a specific inhibitor of CaMKII. The final adenovirus (AdEmpty) served as a control, expressing an empty adenoviral backbone. A map of the three adenoviral vectors, which were purchased from VectorBuilder, USA, are shown in Fig. S1.

The day after seeding, chondrocytes were infected with recombinant adenoviruses at 10, 50 or 100 Multiplicity of Infection (MOIs) as described in the individual experiments in 2% complete medium (2%CM - DMEM F12 (Thermo Fisher, Waltham, MA, USA), 2% Fetal Bovine Serum (FBS; Gibco Invitrogen, Carlsbad, CA, USA) and 1% of antibiotics and antimycotic solution (Sigma-Aldrich, St. Louis, MO, USA)-). Twenty-four hours following infection, viral inoculums were removed and replaced with 2%CM.

### 2.3 CaMKII Assay

To measure CaMKII activity, 4.5 × 10^4 AHACs were plated in 24-well plates and cultured in 10% complete media 10%CM for 24 hours. Following this, they were infected at an MOI of 100 with either AdCaMKII or AdEMPTY in 2% CM for 48 hours. Cell lysates were generated by lysing the chondrocytes in RIPA buffer (50 mM Tris, pH 8; 150 mM NaCl; 1% Triton X-100; 0.1% SDS; 0.5% sodium deoxycholate; 10% protease inhibitor; 1% phosphatase inhibitor, both from Sigma-Aldrich, St. Louis, MO, USA). The lysates were centrifuged at 15,000 G for 20 minutes at 4°C, and the supernatant was stored at -80°C until use in the kinase assay. CaMKII activity was tested with the SignaTECT™ kinase kit (Promega) according to the manufacturer’s instructions. CaMKII activity was quantified following densitometric analysis of the spots on the control membrane – which measured the background levels of CaMKII activity between individual samples.

### 2.4 RT-PCR analysis

For gene expression analysis, 4.5×10^4^ AHACs were plated in 24 well plates and cultured in CM for 24 hours. The cells were washed in PBS and treated as described in the results section. Total RNA was extracted using TRIzol reagent (Invitrogen, Carlsbad, CA, USA), according to the manufacturer’s instructions. A minimum of 250ng of RNA was reverse transcribed to cDNA using a High-Capacity cDNA Reverse Transcription Kit using random primers (Applied Biosystems, Waltham, MA, USA). Quantitative PCR was performed with hot-start DNA polymerase (Qiagen, Hilden, Germany), as previously described (9). Primer sequences are listed in Supplementary Table 1 (All primers were purchased from Sigma-Aldrich, St. Louis, MO, USA). All data were normalised to internal control of 18S values. All experiments were performed in a PCR system (CFX Opus 384 Real-Time PCR System; BioRad, Hercules, CA, USA).

### 2.5 Micromass Cultures

AHACS’s were plated at a density of 1×10^7^ cells/mL in 20µl of 10%CM in 24-well plates. Cells were left to attach for 3h and then 500µl of 2%CM was added. Upon different treatment, the micromasses (MM) were stained with alcian blue to measure proteoglycan content, as previously described (12).

The optical density of the extracted dye was read at 630nm with a ClarioStar spectrophotometer (BMG LABTECH, Offenburg, Germany). Absorbance values were normalised to protein content of respective samples as quantified using the bicinchoninic acid (BCA) assay, following manufacturer instructions (ThermoFisher Scientific; Waltham, MA, USA). Images of the micromasses were acquired at room temperature with a stereomicroscope (SZTL 350 Stereo Binocular Microscope, VWR®; Radnor, PA, USA).

### 2.6 Immunofluorescence assays

For immunofluorescent assays, chondrocytes were seeded on Lab-Tek chamber slides and infected with recombinant adenoviruses for 48 hours. Cells were washed once in PBS, then fixed in 4% buffered Paraformaldehyde (PFA), pH 7.4 for 15 mins at room temperature. After three washes with PBS, autofluorescence was quenched with 50mM Ammonium Chloride for 10 minutes. Cells were then permeabilised with 0.1% Triton X diluted in PBS for 5 minutes. A blocking buffer (0.5% BSA, 2% goat serum in PBS) was then added to the cells for 1 hour at room temperature. Cells were incubated overnight at 4°C with anti-Metalloprotease 13 (MMP13) mouse monoclonal antibody (diluted 1:100 in blocking buffer; Santa-Cruz), anti-Ki67 rabbit monoclonal antibody (diluted 1:100 in blocking buffer; Santa-Cruz), or in blocking buffer alone as a negative control. After washing with TBS-0.1% tween three times, cells were incubated with Alexa 647 conjugated goat anti-mouse IgG secondary antibody (diluted 1:200 in blocking buffer; Abcam) for one hour at room temperature. After a further three washes in TBS-0.1% tween, cells were incubated with 18µM DAPI in PBS for five minutes. Following a final three washes in TBS-0.1% tween, slides were mounted in Mowiol.

Images were acquired using a confocal microscope (Nikon A1M Confocal Microscope) using a 40x objective lens. After acquisition, images were enhanced for best graphic rendering using Nikon Elements, all with the same parameters, including the control image. The fluorescence intensity per cell was measured using Cell Profiler Image Analysis Software (13). The whole cell MMP13 expression as shown in Fig, 1 d, were generated using a minimum of 100 cells, per condition, per biological donor.

**Figure 1.**
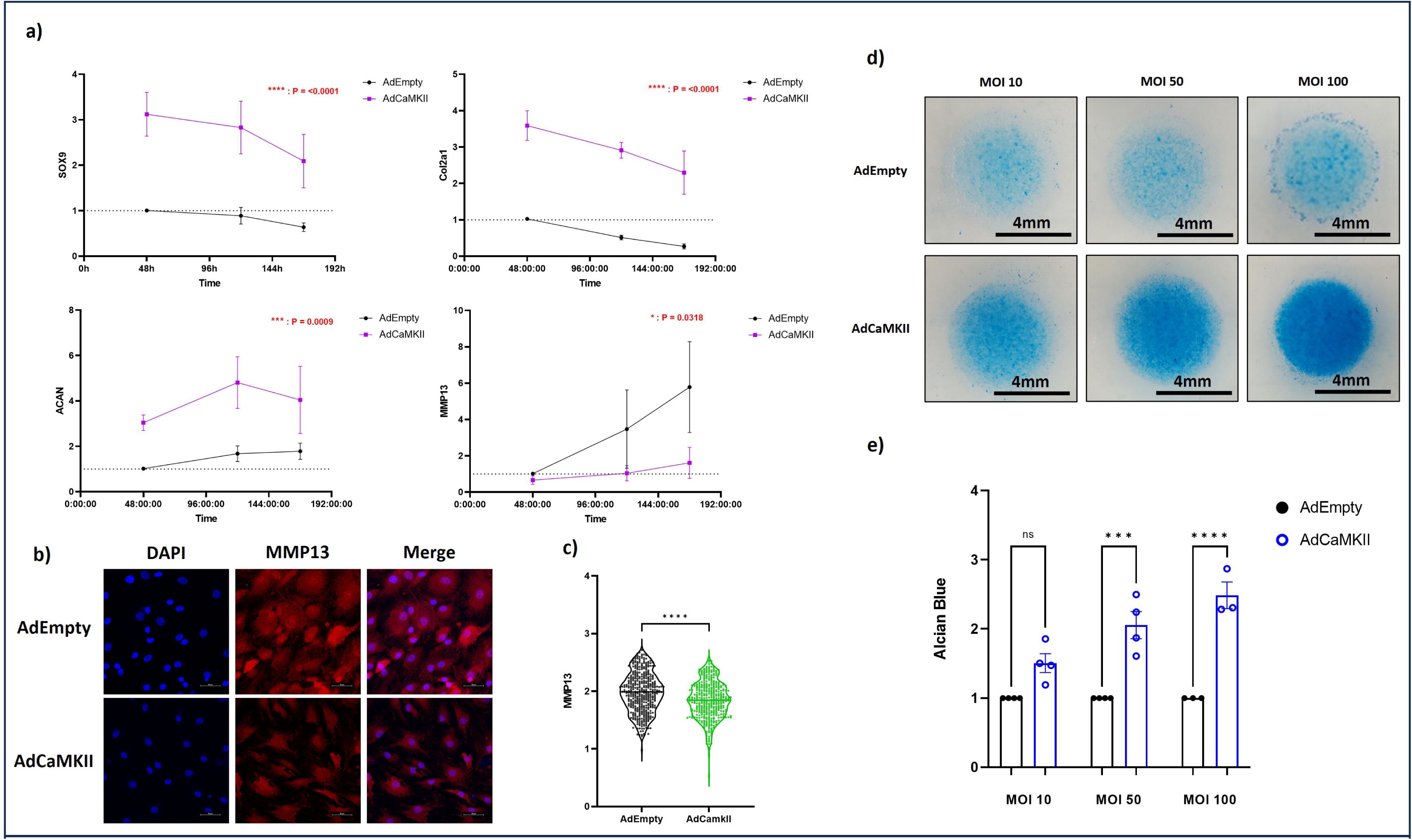
CaMK/1 Overexpression Promotes Anabolic Activity and Reduces Expression of Catabolic Enzymes in AHACs. **a)** *SOX9, Co/2al, ACAN* and *MMP13* mRNA expression in AHACs infected with either AdCaMKII or AdEMPTY (MOl 100) over a 7-day time course. N=5. Data were analysed with a 2way ANOVA. **b)** Representative images of MMP13 expression in AHACs infected with AdCaMKII or AdEMPTY (MOl 100) for 48hours. **c)** Quantification of expression of MMP13 in AHACs following infection with AdCaMKII or AdEMPTY for 48hours (MOl 100). N=3. Data were analysed with a Mann-Whitney Test(****= P<0.0001). **d)** Representative images of micromass cultures stained with Alcian Blue after infection with MOll0, 50 or 100 of AdCaMKII or AdEMPTY over a 5-day period. **e)** Quantification of Alcian Blue intensity following extraction with Guanidine hydrochloride. N=4 for MO ls 10 and 50. N=3 for MOl 100. Data were analysed with a one-way ANOVA (ns = not significant,***= P<0.0005, **** = P<0.0001).

### 2.7 SDS-PAGE and Western Blotting

After treatments, cells were washed once in cold PBS. Protein was extracted by scraping cells in RIPA buffer and leaving the lysates on ice for 30minutes with periodic vortexing. Cell debris was removed by centrifuging lysates at 15,000G for 20 minutes at 4°C. Protein concentration was determined by BCA assay (Thermo Fisher, Waltham, MA, USA) according to manufacturer instructions.

Protein extracts were resolved by SDS-PAGE, then transferred to a nitrocellulose membrane (Cytivia Amersham; Amersham, Buckinghamshire, UK) using a wet transfer system (BioRad; Hercules, CA, USA) at 100V for 60 minutes. P62 was detected with anti-P62 mouse monoclonal (diluted 1:1,000; Santa Cruz, California, US), phospho-Beclin1 (pBeclin1) (Ser90/Ser93/Ser96) was detected with anti-pBeclin1 (diluted 1:1,000; Affinity Biosciences, Melbourne, Victoria, AU) rabbit polyclonal antibody and Beclin 1 was detected with anti-Beclin 1 (diluted 1:1,000; Affinity Biosciences, Melbourne, Victoria, AU) rabbit polyclonal antibody. All were then detected using HRP-conjugated secondary antibodies (diluted 1:10,000, Abcam, Cambridge, UK) and the Supersignal^TM^ West Pico PLUS chemiluminescent substrate (Thermo Fisher, Waltham, MA, USA). Glyceraldehyde 3-phosphate dehydrogenase (GAPDH) was detected with anti-GAPDH (diluted 1:2,000; Abcam, Cambridge, UK) rabbit polyclonal antibody, followed by incubation with anti-Rabbit Alexa Flour^TM^ 680 conjugated secondary antibody (diluted 1:10,000; Abcam, Cambridge, UK). For pBeclin1 analysis, the membranes were blocked in 5%BSA/0.01%tween 20 in TBS. For Beclin 1 analysis, the membranes were blocked in 5% milk powder/0.01%tween 20 in TBS. For all other antibodies the membranes were blocked in protein free blocking buffer in TBS (Thermo Fisher, Waltham, MA, USA). Primary antibodies were diluted with respective blocking buffers.

### 2.8 Statistical Analysis

Shapiro-Wilk tests were carried out to assess data for normality. Parametric data between two groups were compared with a t-test. When the standard deviations (SD’s) between groups could not be assumed to be equal a Welch’s correction was applied. Where there were multiple groups, we used analysis of variance (ANOVA) analysis, with a Bonferroni post-test to correct for multiple comparisons using statistical hypothesis testing. Where SD’s could not be assumed to be equal, a Browne-Forsythe and Welch ANOVA test was applied with a Dunnett T3 post-test. Where there were multiple groups and multiple time points, we used a two-way ANOVA with a Bonferroni post-test. Nonparametric data between two groups were analyzed by a Mann-Whitney test. Where there were multiple groups, a Kruskal-Wallis test with a Dunn’s multiple comparisons test was applied. P-values <0.05 were considered significant. *, P < 0.05; **, P < 0.005; ***, P < 0.0005; ****, P < 0.0001. All the data in the graphs are expressed as mean ± SEM. All statistical analyses were performed using GraphPad Prism version 9.4.1 for Windows. (GraphPad Software, Boston, Massachusetts, USA)

## 3 Results

### 3.1 Overexpression of constitutively activated CaMKIIγ promoted anabolism and decreased expression of catabolic enzymes in human articular chondrocytes

To better understand how CaMKII modulates metabolism in articular chondrocytes, we generated an adenovirus expressing a constitutively active form of CaMKIIγ (AdCaMKII, Fig. S1C). Expression of the transgenes were validated by qPCR (Fig. S2A,B). To confirm activity of the recombinant CaMKIIγ, we infected AHACs in monolayer at MOI 100 for 48 hours and quantified CaMKII activation by measuring phosphorylation of a biotinylated CaMKII substrate with phosphorous-32 (^32^P) using the SignaTECT Calcium/Calmodulin-Dependant Protein Kinase Assay System (Fig. S2C). To confirm the activity of AIP delivered through the virus, we compared the expression levels of HMOX1, a selective transcriptional target of CaMKII inhibition (2), in AHACs infected at MOI 100 with AdAIP or stimulated with 5μM of soluble AIP. Upregulation of HMOX1 in response to the virus or to the soluble inhibitor was comparable (Fig. S2D).

To test the effect of CaMKII on chondrocyte differentiation, we then infected AHACs with AdCaMKII or with the AdEmpty control and measured the expression of chondrocyte phenotypic markers by qPCR over a seven-day time course. Overexpression of activated CaMKII increased the expression of SRY-box transcription factor 9 (*SOX9*), Collagen type II (*COL2A1*) and Aggrecan (*ACAN*) mRNAs and decreased the expression of *MMP-13* at all the time points in comparison to cells infected with the control virus (Fig. 1A).

Downregulation of MMP-13 was confirmed at protein level in AHACs infected with AdCaMKII or AdEmpty for 48 hours by immunofluorescence (Fig. 1B, C).

To confirm the pro-anabolic effect of activated CaMKIIγ, we measured proteoglycan content in AHAC micromass cultures infected with AdCAMKII or AdEmpty at different MOIs. Overexpression of constitutively activated CaMKII increased GAG content in a dose-dependent manner (Fig. 1D, E).

### 3.2 CaMKII counteracts the catabolic effect of IL-1β in AHACs

Interleukin 1-β (IL1-β) is a pro-inflammatory cytokine whose expression in the AC increases during the progression of OA (14,15). To investigate whether the overexpression of activated CaMKIIγ could counteract the pro-catabolic effect of IL-1-β, we infected AHACs with AdCaMKII or AdEmpty for four days and then added recombinant human IL1-β (20ng/mL) for 24 hours.

Overexpression of activated CaMKII avoided a decrease in proteoglycan content in AHAC micromasses exposed to IL1-β for 24 hours (Fig. 2A, B).

**Figure 2.**
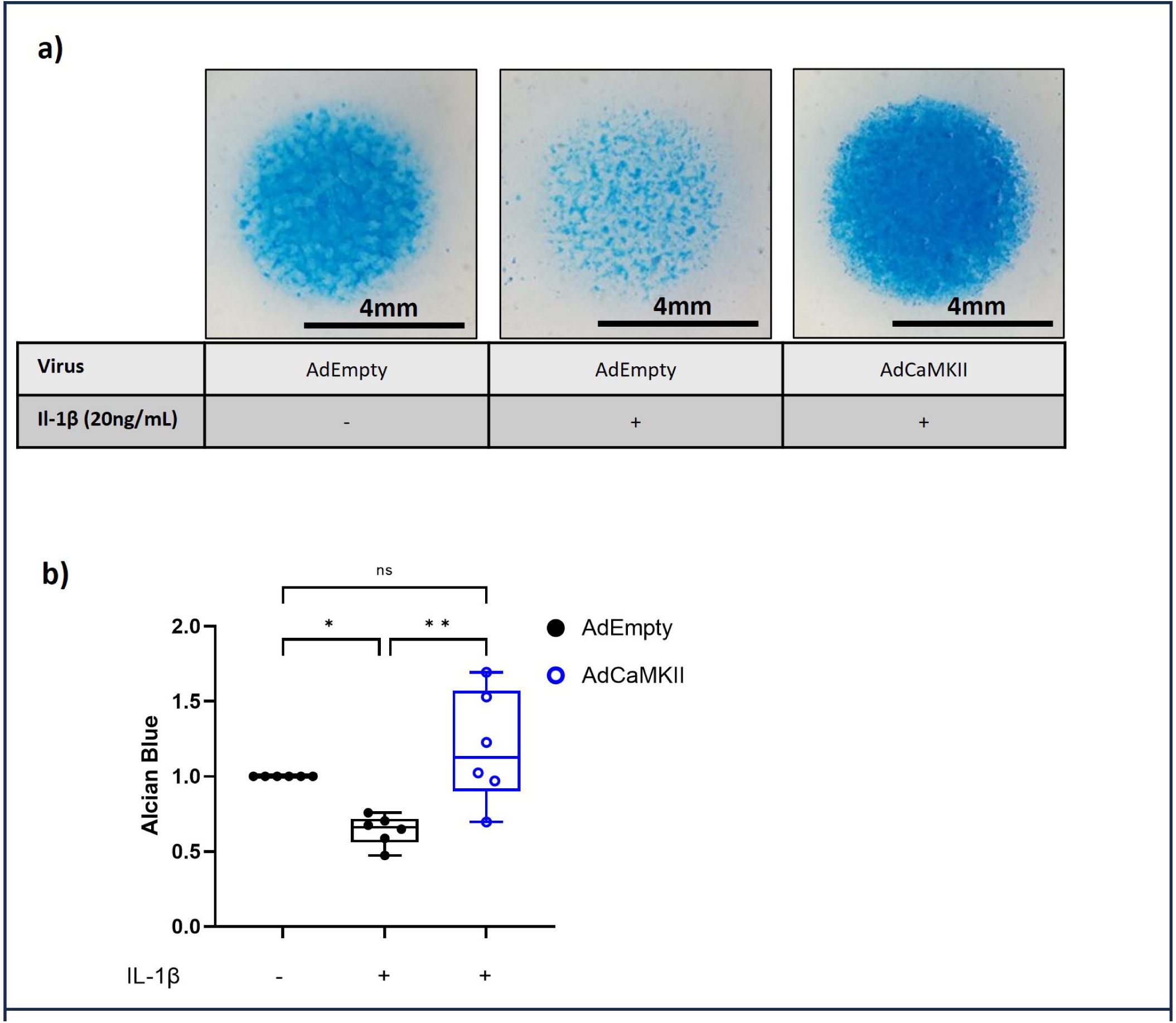
CaMK/1 overexpression prevents inflammation mediated decreases in anabolic activity. **a)** Representative images of micromass cultures stained with Alcian Blue after infection with AdCaMKII or the AdEMPTY (MOI 100) over a 5-day period. Respective wells were treated with 20ng/ml ILlP for 24 hours. **c)** Quantification of Alcian Blue intensity following extraction with Guanidine hydrochloride. N=6. Data were analysed with a one-way repeated measures ANOVA (* = P<0.05, ** = P<0.005).

Overall, our results suggest a pro-anabolic role of CaMKII in the articular chondrocytes and an important homeostatic function of the kinase in counteracting catabolic stimuli upregulated in OA such as IL-1-β.

### 3.3 Overexpression of CaMKII decreases proliferation and increases hypertrophic markers

Previous studies have shown that CaMKII has a role in the transition from proliferating chondrocytes to pre-hypertrophic, then hypertrophic chondrocytes in the growth plate during bone growth (4,8,16). Furthermore, Saitta and colleagues showed the upregulation of hypertrophic markers in response to Bone Morphogenetic 4 (BMP4) is mediated by CaMKII in human fetal chondrocytes (3).

We observed changes in the morphology and viability of AHACs overexpressing AdCaMKII in comparison to cells infected with the control virus, as cells acquired a round rather than an elongated shape (FIG. 3A). We therefore carried out qPCR analysis of hypertrophic marker, Type X Collagen (*TXCOL)*, over a 7-day time-period in chondrocytes overexpressing activated CaMKII. Confirming results in other biological contexts, we observed increased expression of *TXCOL* over time in AHACs infected with AdCaMKII. Furthermore, overexpression of activated CaMKII for 48h induced a significant reduction of the percentage of proliferating chondrocytes (Fig. 3C, D), as shown by decreased Ki67 staining. This indicates that CaMKII’s role in chondrocyte maturation, observed across various chondrocyte populations, is conserved in AHACs.

**Figure 3.**
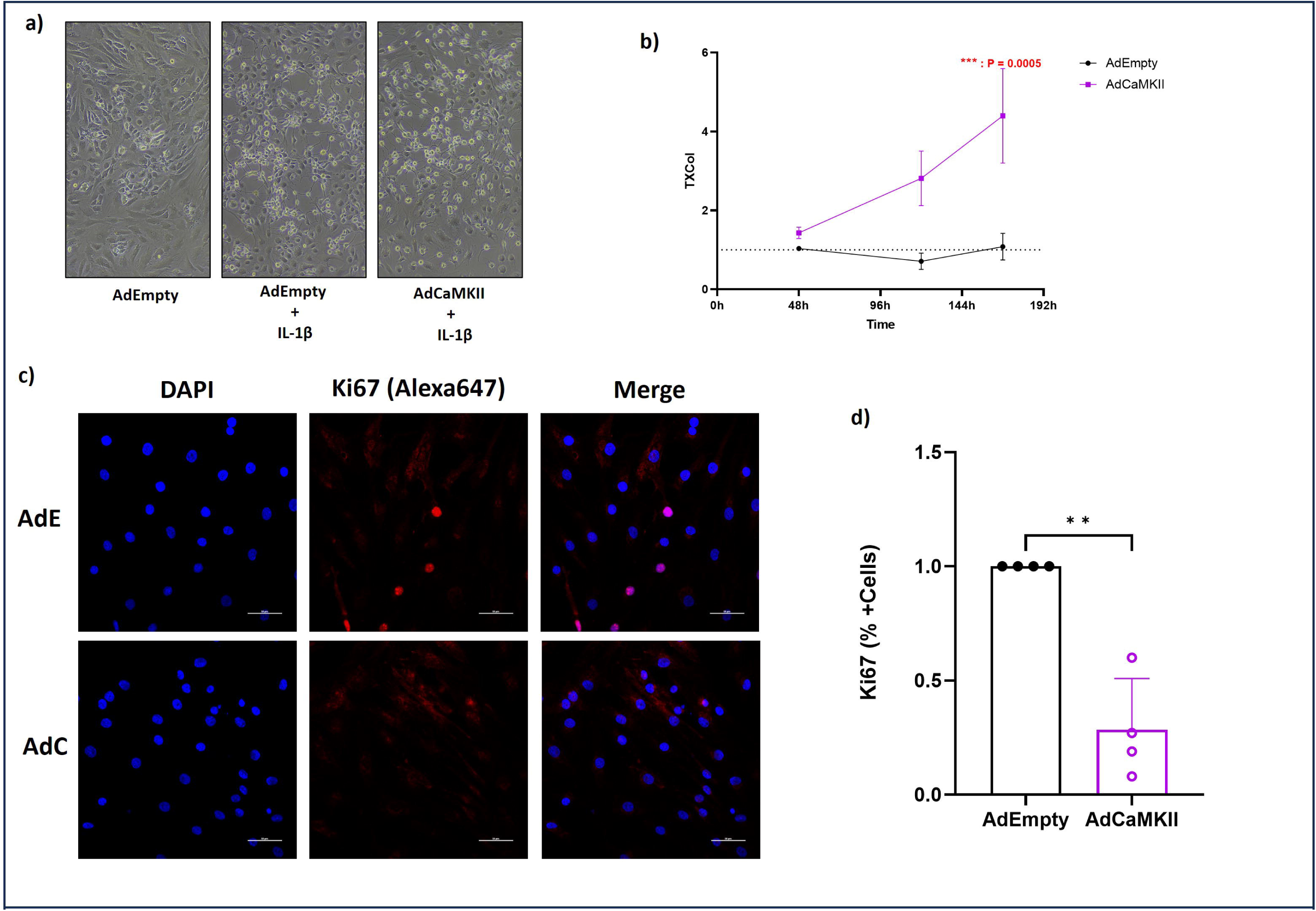
Overexpression of CaMK/1 inhibits proliferation and induces hypertrophy in Articular Chondrocytes. **a)** Representative brightfield images of AHACs infected with AdCaMKII or AdEmpty (MOI 100) for Sdays in the presence/absence of Ill (20ng/mL) for 24hours. **b)** TXCol mRNA expression in AHACs infected with AdCaMKII or AdEMPTY (MOI 100) over a 7-day time course. N=S. Data were analysed with a 2way ANOVA. c) Representative images of Ki67 staining in AHACs infected with either AdEmpty or AdCaMKII (MOl 100) for 48 hours. **d)** Percentage of AHACs positively stained with Ki67 after infection with AdCAMKII or AdEMPTY (MOl 100) for 48 hours. N=4. Data were analysed with a paired T-Test (** = P<0.005).

### 3.4 CaMKII Induces Autophagy in Articular Chondrocytes

Autophagy is a process which has been associated with chondrogenic differentiation (17). Autophagy plays a protective role in maintaining chondrocyte health and function through providing energy in low nutrient environments such as the hypoxic conditions of the AC (18). Furthermore, recent work suggested that autophagy is involved in preventing premature hypertrophic differentiation during development (19).

To gain further insight on how CaMKII is modulating chondrocyte phenotype and metabolism, we stimulated AHACs with AdCaMKII, AdAIP or AdEmpty and quantified the expression of phospho-Beclin1, which is required for autophagosome formation (20). Indeed, infection of chondrocytes with AdCaMKII not only promoted phosphorylation of Beclin1 (Fig. 4A, B) but also decreased the expression of the ubiquitin binding protein (p62). P62 is a ubiquitin sensing protein which transports proteins to the autophagosome and accumulates upon defective autophagy (21). Inhibition of CaMKII in AHACs infected with AdAIP increased the expression of p62 in comparison to cells infected with AdEmpty (Fig4C, D). These results overall point to CaMKII as a modulator of autophagy in the AC.

**Figure 4.**
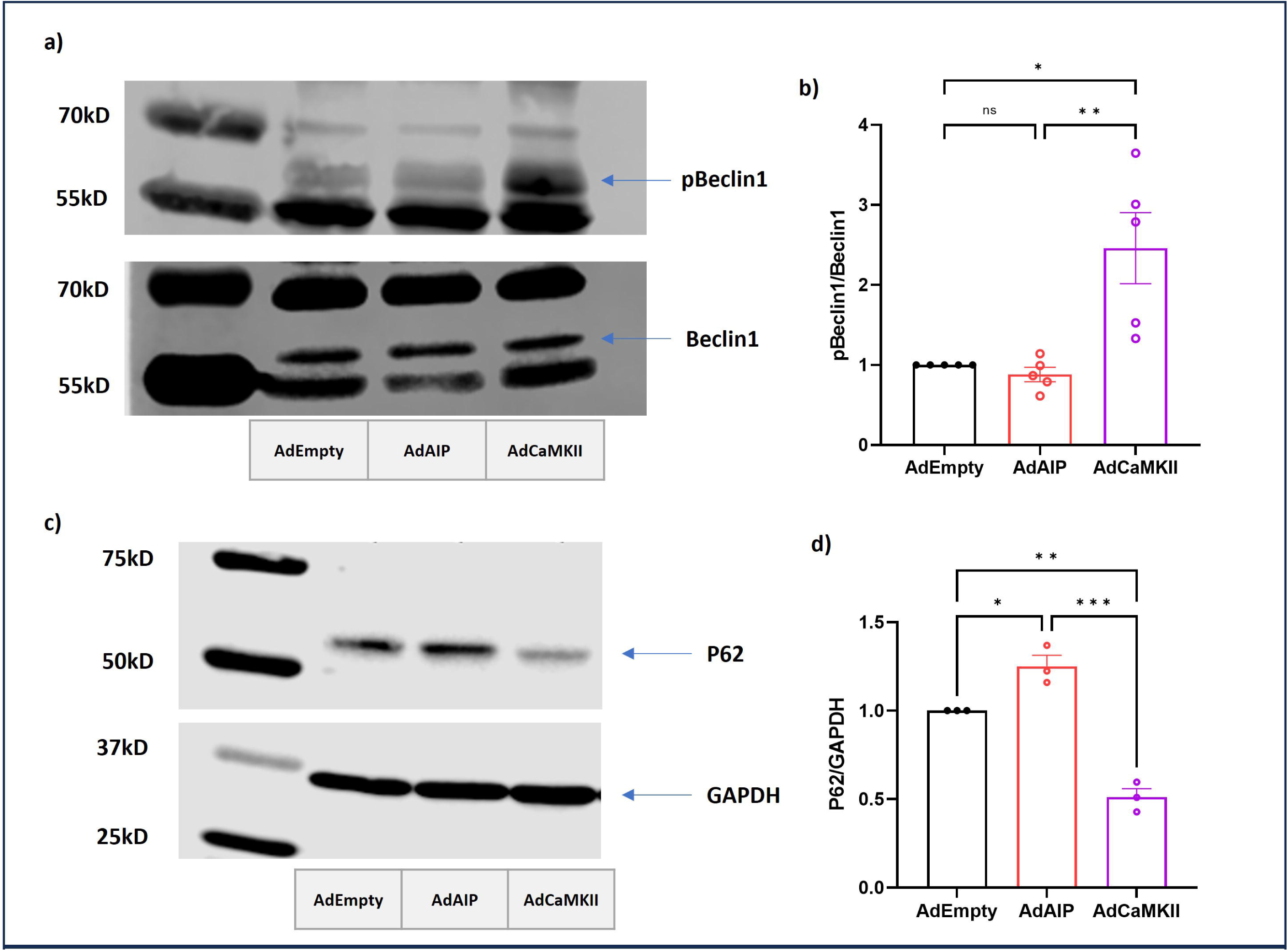
CaMK/1 induces Autophagy in Articular Chondrocytes. **a)** Representative image of a Western blot for phosphorylated/total Beclinl following infection of AHACs with MOI 100 AdEmpty, AdalP or AdCaMKII for 48 hours. **b)** Densitometric analysis of Western blot for pBeclinl and Beclinl. Bars represent ratio of pBeclinl/Beclinl +/-SEM. N=S. Data were analysed with a one-way repeated measures ANOVA (* = P<0.05, ** = P<0.005). c) Representative image of a Western blot for P62 in AHACs infected with MOI 100 AdEmpty, AdalP or AdCaMKII for 48 hours **d)** Densitometric analysis of Western blot for P62. Bars represent P62 expression normalised to GAPDH loading control +/-SEM. N=3. Data were analysed with a one-way repeated measures ANOVA (* = P<0.05, ** = P<0.005, *** = P<0.0005)

### 3.5 The CaMKII-dependant Anabolic Response is prevented upon inhibition of Autophagy

Autophagy is vital for maintaining energy metabolism in the body. Studies have shown that diabetic patients are at higher risk of cartilage damage if autophagy decreases (22), as well as decreases in expression of genes involved in autophagy in osteoarthritic chondrocytes (23). For this reason, we hypothesised that the increases in anabolic activity of AHACS’s following overexpression of activated CaMKII are dependent on energy and metabolites produced through autophagy. To test this, we overexpressed activated CaMKII in the presence of 5nM Bafilomycin, an inhibitor of autophagy (24), over a 48-hour period and quantified the production of GAG’s in micromass cultures. The increase in GAG’s induced by CaMKII, were completely recovered to baseline levels upon the inhibition of autophagy (Fig. 5A, B), suggesting that the anabolic activity induced by CaMKII is dependent on autophagy.

**Figure 5.**
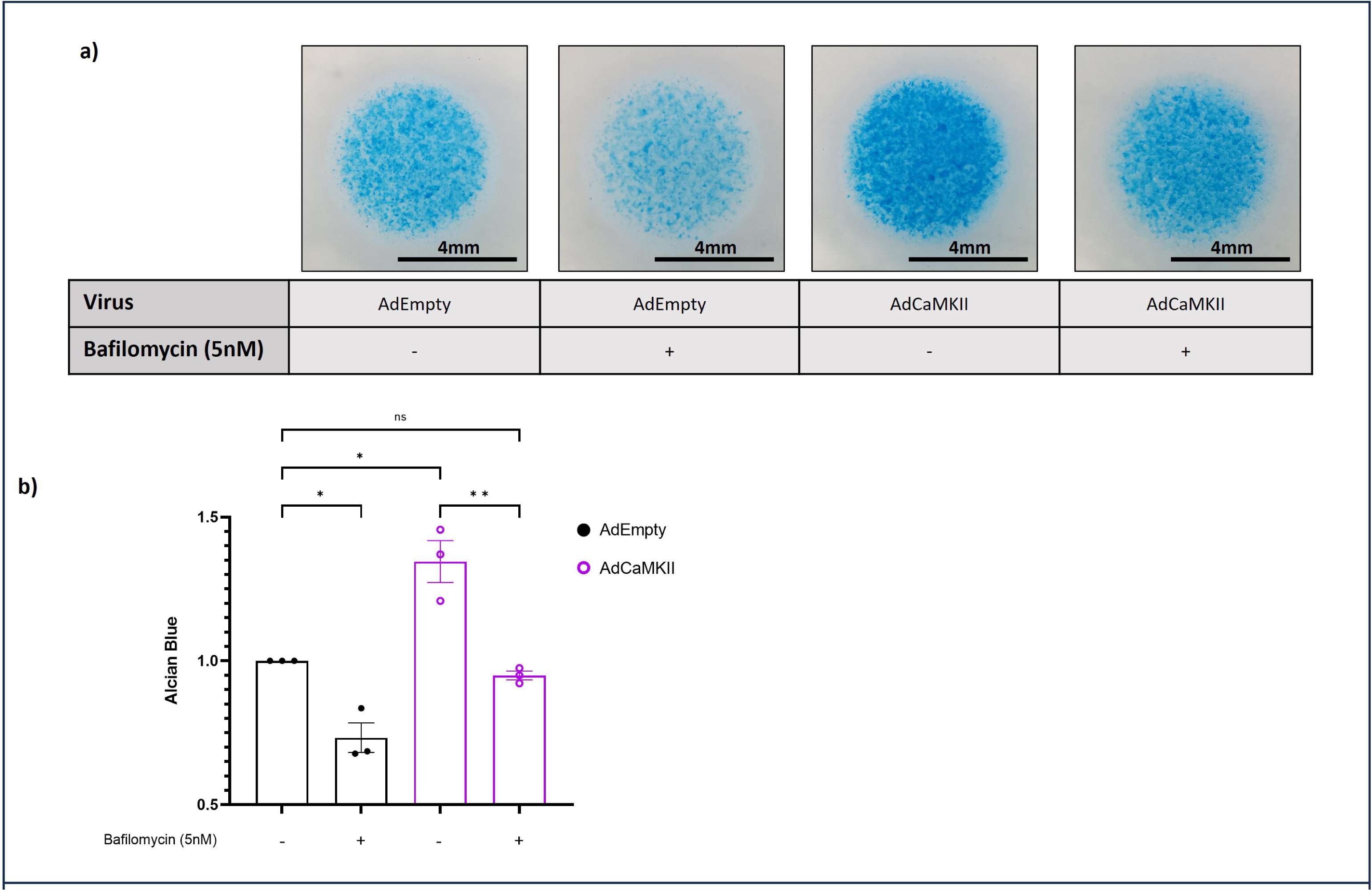
Inhibition of Autophagy prevents the CaMK/1 induced anabolic response. **a)** Representative images of AHAC micromass cultures stained with Alcian Blue after infection with AdCaMKII or AdEMPTY for 48hours and co-stimulated with SmM bafilomycin for 24 hours. b) Quantification of Alcian Blue intensity following extraction with Guanidine hydrochloride. N=3. Data were analysed with a one-way repeated measures ANOVA (* = P<0.05, ** = P<0.005).

## 4 Discussion

Here we characterised the role of CaMKII in the regulation of features of chondrocyte metabolism and showed its involvement in the modulation of hypertrophic differentiation as well as autophagy in AHACs. This study builds on our previous work, as well as that from other groups, which indicated an important homeostatic role for this kinase both in vitro and in vivo. (2–5,8).

CaMKII has been extensively explored in other biological systems. It has a crucial function in the brain where it is required for synaptic transmission and memory formation (25). CaMKII is also the most abundant calcium calmodulin kinase in the heart where it regulates the excitation-contraction coupling (26). Overactivation of CaMKII has been linked to myocardial disease, as it can promote apoptosis, arrhythmias (27) and pathological hypertrophy (28).

CaMKII activation has also been associated with hypertrophic maturation of chondrocytes during development. Retroviral mediated constitutive expression of CaMKII in the developing chicken wing promoted premature chondrocyte differentiation and elongation of skeletal elements (4). Similarly, in mouse, downregulation of CaMKII activity halted the transition from proliferating to hypertrophic chondrocytes during bone development (29).

Activated CaMKII is upregulated in the articular cartilage during OA progression both in human and mouse (2). Sugita et al showed that the expression of CaMKIIδ increases in a murine model of OA, and that the kinase can form a protein complex with the cofactor Hairy and Enhancer of Split-1 (HES1), transforming it from a transcriptional repressor into a transcriptional activator and promoting the expression of catabolic genes, including MMP13 and IL6 (5).

Considering that re-start of the developmental programme and hypertrophic maturation of articular chondrocytes is associated with AC degeneration in OA (30,31), we previously considered inhibition of CaMKII as a potential strategy to halt catabolism in the AC in OA patients. However, we showed that pharmacological inhibition of CaMKII with KN93 exacerbated cartilage degeneration in a surgically induced mouse model of OA. Inhibition of CaMKII acted in synergy with IL-1β in promoting a catabolic response on isolated chondrocytes (2).

Our gain of function experiments helped in unravelling the complexity of the CaMKII-mediated signalling in the AC and placed the molecule at the crossroads between multiple homeostatic functions.

Our data described an important anabolic function for CaMKII, as its overactivation promoted synthesis of ECM components and prevented IL-1β-mediated proteoglycan degradation in chondrocytes. Active CaMKIIγ also induced downregulation of MMP13 expression, a metalloproteinase promoting collagen degradation and ECM destruction in OA (32).

Furthermore, here we showed for the first time that activation of autophagy is also involved, at least in part, in promoting anabolism in response to CaMKII activation. Activation of autophagy induces the degradation of damaged or faulty organelles and proteins which otherwise cause aggregates, leading to cellular dysfunction or apoptosis (33). Studies conducted by Lotz’s group and others have highlighted the crucial role of autophagy in maintaining cartilage homeostasis. They have shown that the decline of autophagy with aging is linked to the development and progression of osteoarthritis (34–36). The involvement of CaMKII in regulating autophagy has also been described in different biological systems (37).

However, while promoting anabolism, we also observed that overexpression of activated CaMKII halted cell proliferation and promoted expression of Collagen Type X, confirming the involvement of the kinase in hypertrophic maturation, as observed during bone development (4,8).

Interestingly, recent literature suggested a link between autophagy and hypertrophic differentiation. Knock-out of ATG13, a subunit of the autophagic initiation complex, led to inhibition of autophagy and acceleration of chondrocyte maturation in zebrafish (19). Sulphation of proteoglycans is also linked to activation of autophagy: knock down of the Sulphatase-Modifying Factor 1 (SUMF1) in mice led to an increase in the number of autophagosomes as a result of impaired lysosomal function, ultimately leading to inhibition of autophagy and resulting in cell death. Interestingly, deletion of SUMF1 resulted in decreased chondrocyte proliferation and affected their maturation, linking therefore modulation of autophagy with chondrocyte maturation during skeletal development (38).

Our data, alongside previous work, confirm the complexity of the CaMKII-mediated signalling and how its careful dissection could lead to the development of new strategies aimed to specifically stimulate its anabolic potential without inducing activation of pro-degenerative mechanisms. In a recent review, Rostas and Skelding summarised our current knowledge on how cellular microenvironment can influence how CaMKII is activated and can lead to different biological responses (39). This, however, is only one of the layers of complexity in the modulation of CaMKII activity. Its expression in multiple isoforms, isoform variants (eg. 11 for CaMKIIδ alone), additional subtypes of CaMKII, holoenzyme confirmations made of multiple isoforms in different proportions within cells and tissues (40) and its subcellular localisation either anchored to the cell membrane, within the cytoplasm (9), in the mitochondrial membrane (41) and within the nucleus (9), are all elements which add to the complexity of the modus operandi of this kinase and that will be worthy of exploring, to fully understand the role of this kinase in the maintenance of cartilage homeostasis, and to understand how to harness it for therapeutic purposes.

## Author Contributions

GN conceived and designed the study. GN and BF supervised the study. NJD conducted the investigation. DS and VA provided the patient samples. NJD, AD and CSH obtained the patient samples. NJD performed the analysis. NJD and GN wrote the original draft of the manuscript. GN, BF, DS, VA, AD and CSH reviewed and edited the manuscript.

## Conflict of Interest Statement

The authors declare no conflict of interest.

## Data Availability Statement

All data generated or analyzed during this study are included in this published article and its supporting figures.

## Notes

### Competing Interest Statement

The authors have declared no competing interest.

