## Supplementary figures and images for "CaMKII induces an autophagy-dependent anabolic response in Articular Chondrocytes"

### Supplementary Figure 1

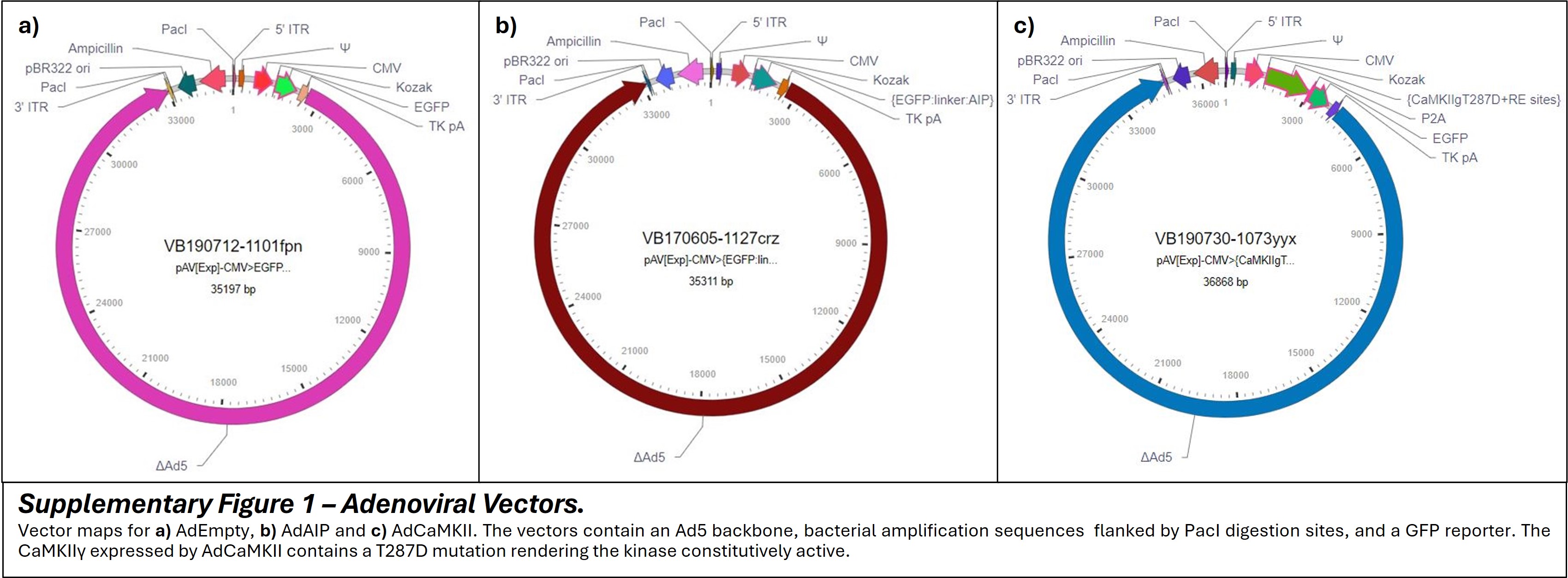

### Supplementary Figure 2

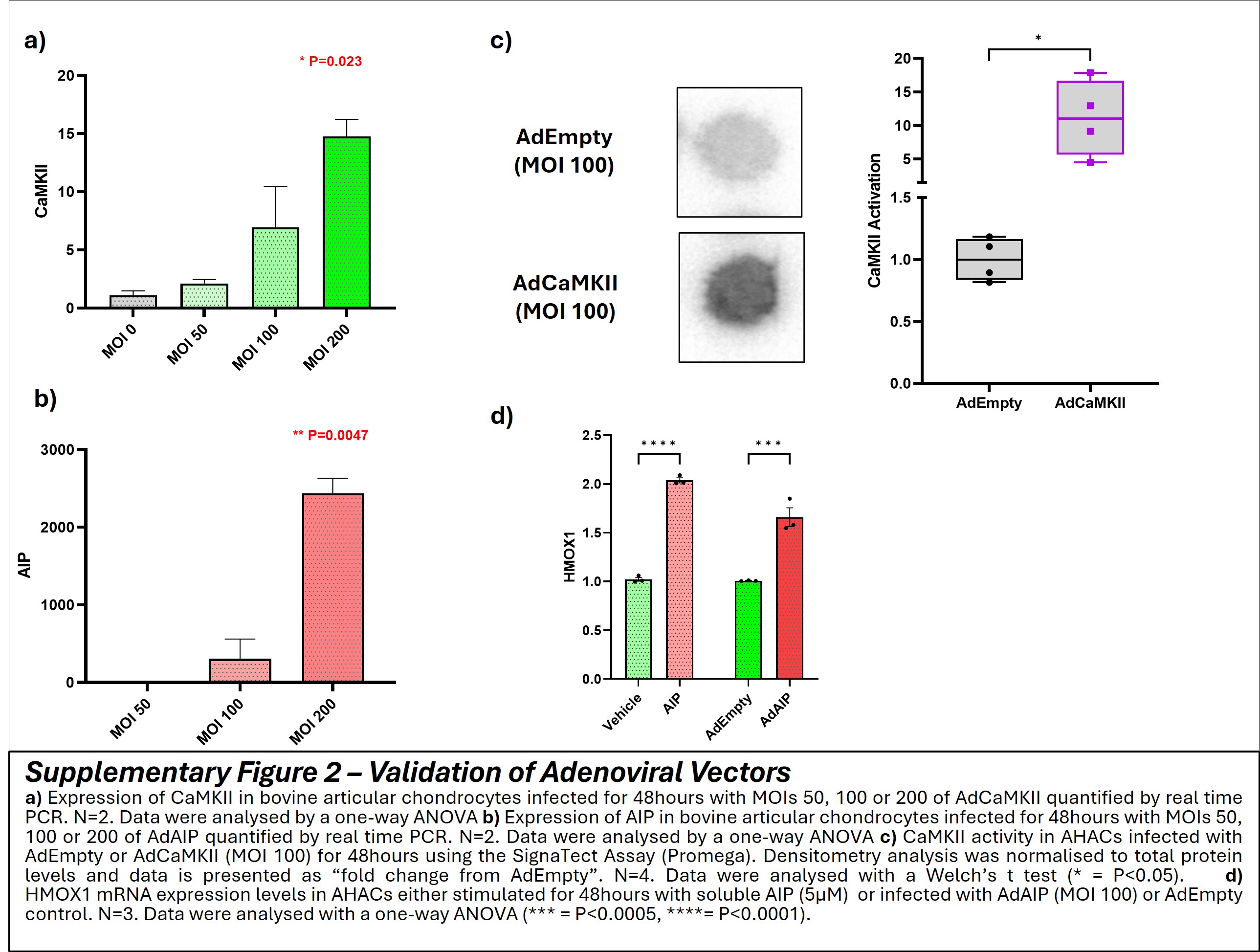

### Supplementary Table 1

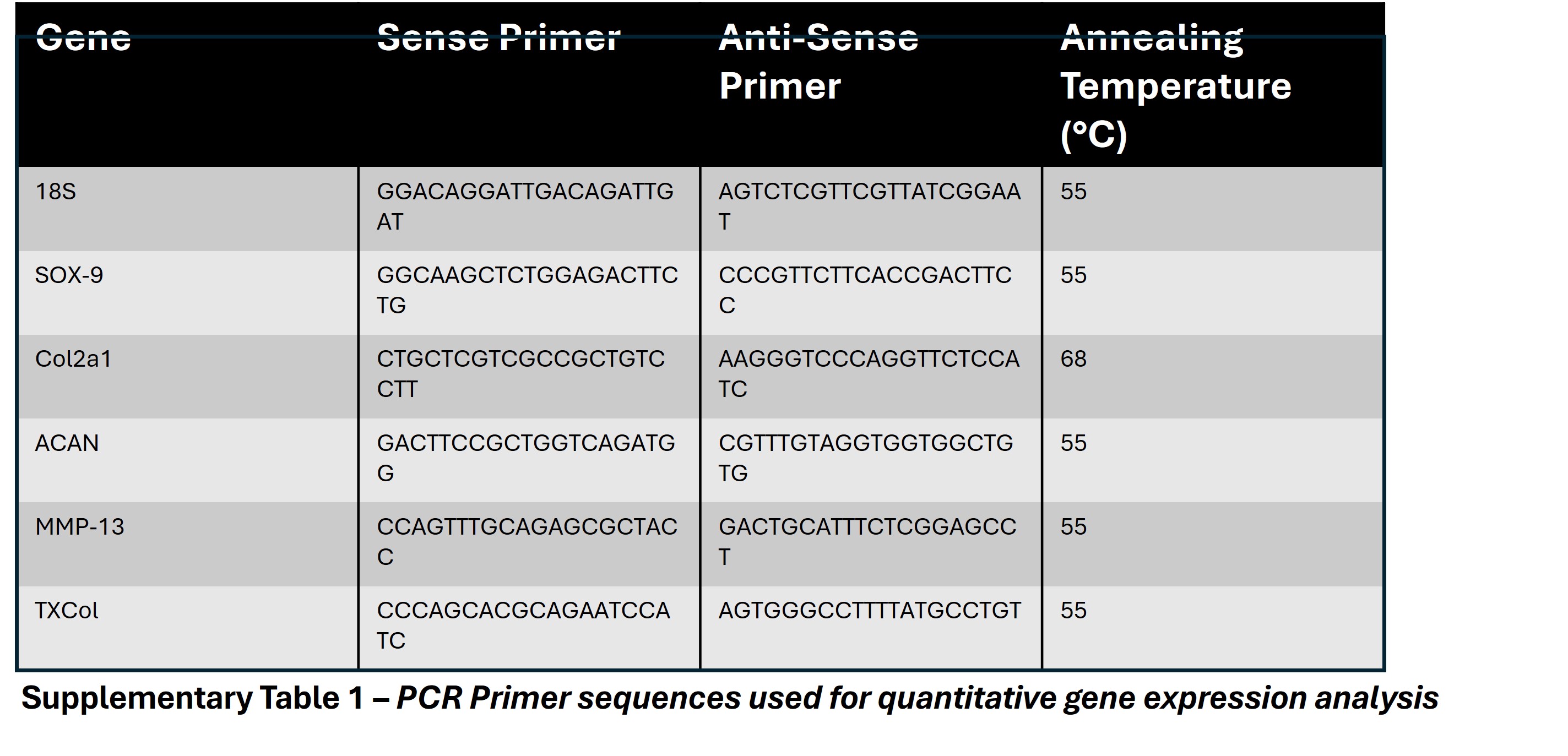
